# The role of liprin-α1 phosphorylation in its liquid-liquid phase separation: regulation by PPP2R5D/PP2A holoenzyme

**DOI:** 10.1101/2024.06.18.599485

**Authors:** Abigail Mayer, Rita Derua, Elijah Spahn, Iris Verbinnen, Yang Zhang, Brian Wadzinski, Mark R. Swingle, Richard Honkanen, Veerle Janssens, Houhui Xia

**Author notes:** Contributed equally (Abigail Mayer, Rita Derua). **Correspondence:** Veerle Janssens; Houhui Xia.

## Abstract

Liprin-α1 is a widely expressed scaffolding protein responsible for regulating cellular processes such as focal adhesion, cell motility, and synaptic transmission. Liprin-α1 interacts with many proteins including ELKS, GIT1, liprin-β, and LAR-family receptor tyrosine protein phosphatase. Through these protein-protein interactions, liprin-α1 assembles large higher-order molecular complexes; however, the regulation of this complex assembly/disassembly is unknown. Liquid-liquid phase separation (LLPS) is a process that concentrates proteins within cellular nano-domains to facilitate efficient spatiotemporal signaling in response to signaling cascades. While there is no report that liprin-α1 spontaneously undergoes LLPS, we found that GFP-liprin-α1 expressed in HEK293 cells occasionally forms droplet-like condensates. MS-based interactomics identified Protein Phosphatase 2A (PP2A)/B56δ (PPP2R5D) trimers as specific interaction partners of liprin-α1 through a canonical Short Linear Interaction Motif (SLiM) in its N-terminal dimerization domain. Mutation of this SLiM nearly abolished PP2A interaction, and resulted in significantly increased LLPS. GFP-liprin-α1 showed significantly increased droplet formation in HEK293 cells devoid of B56δ (PPP2R5D knockout), suggesting that PPP2R5D/PP2A holoenzyme inhibits liprin-α1 LLPS. Guided by reported liprin-α1 Ser/Thr phosphorylation sites, we found liprin-α1 phospho-mimetic mutant at serine 763 (S763E) is sufficient to drive its LLPS. Domain mapping studies of liprin-α1 indicated that the intrinsically disordered region, the N-terminal dimerization domain, and the SAM domains are all necessary for liprin-α1 LLPS. Finally, expression of p.E420K, a human PPP2R5D variant causing Houge-Janssens Syndrome type 1 (also known as Jordan’s Syndrome), significantly compromised suppression of liprin-α1 LLPS. Our work identified B56δ-PP2A holoenzyme as an inhibitor of liprin-α1 LLPS via regulation at multiple phosphorylation sites.

## Introduction

Efficient spatial and temporal cellular signaling requires the regulation of nano-domains containing a high concentration of proteins. Focal adhesions, synaptic active zones and postsynaptic densities are examples of nano-domains that require large protein networks for proper signaling and maintenance. One mechanism by which protein networks are organized into regulated nano-domains is through liquid-liquid phase separation (LLPS), which is postulated to increase protein concentration via formation of membrane-less liquid droplets [1–3]. The phase separation of many proteins including liprin-α3 and SYD-2 has been shown to play essential roles in critical cellular processes including cell motility and synaptic transmission [1, 4–7]. Studies of proteins participating in LLPS suggest the modulation of oligomerization domains, intrinsic disordered regions, and protein-protein interaction domains are significant for driving the phase separation of individual proteins or multi-protein complexes into condensed, dynamic liquid droplets [8]. The dynamic nature of LLPS requires precise regulation to favor the condensed multivalent interactions. Phosphorylation is a ubiquitous mode of regulation of protein function and has been shown to promote LLPS for scaffolding proteins such as liprin-α3 [1] and SYD-2 [5], the *Caenorhabditis elegans* homologue for mammalian liprin-α (1-4). PKC and SAD-1 kinase activation increase LLPS of liprin-α3 and SYD-2, respectively [1, 5], but little is known about the role protein phosphatases play in regulating liprin-α LLPS.

Liprins (α1-4) are major scaffolding proteins playing significant roles in synapse formation and focal adhesions. While liprin-α1 is ubiquitously expressed in eukaryotic cells, liprin-α2 and liprin-α3 are mainly expressed in neuronal cells [9]. Liprin-α binding to ELKS is important for concentrating active zone assembly proteins and synaptic vesicle recruitment in neurons [10, 11]. Liprin-α1 also localizes to dendritic spines [12] and interacts with GRIP [12], GIT1 [13], and/or LAR-RPTP [14] to regulate AMPA receptor targeting. In non-neuronal cells, liprin-α1 expression promotes cell spreading and the disassembly of focal adhesions [2]. Studies using MDA-MB231 cell lines found liprin-α1 expression increased cell invasiveness, likely via promoting integrin endocytosis and degradation of extracellular matrix [9]. The role of liprin-α1 in cell spreading and migration is regulated via phosphorylation [15, 16], however the phosphorylation sites are not characterized. While liprin-α1 has ubiquitous expression in eukaryotic cells and plays important roles in both neuronal and non-neuronal cells, it is unknown whether liprin-α1 can undergo phosphorylation dependent LLPS.

Several studies identified liprin-α isoforms that undergo LLPS in response to kinase activation; however, there is little known about the phosphatases involved in this process. PP2A is a ubiquitously expressed phospho-Ser/Thr phosphatase complexing with one of the many B subunits, and one of several scaffolding A subunit forming different PP2A holoenzyme for dephosphorylation catalysis. In this study, we aimed to determine whether liprin-α1 could undergo LLPS and the role and identify of PP2A holoenzyme in regulating LLPS. We and others have previously shown liprin-α1 binds to B56δ and B56γ regulatory subunits of PP2A [16–19]; In our current report, we mapped liprin-α1 binding to B56δ through a short linear motif (SLiM) on liprin-α1. Additionally, HEK293 cells transfected with B56δ-binding-deficient liprin-α1 mutants exhibited significantly increased droplet formation. Moreover, B56δ null HEK293 cells displayed an increase in GFP-liprin-α1 LLPS, further implicating B56δ-mediated phosphorylation in LLPS. GFP-liprin-α1 mutant mimicking phosphorylation at one candidate B56δ-PP2A phosphosite (S763) showed increased LLPS. Domain-mapping also indicated the dimerization domain, intrinsic disordered domain, and sterile alpha helical motifs (SAMs) of liprin-α1 are required for LLPS. Interestingly, GFP-liprin-α1 also shows significant LLPS in homozygous HEK293 cells expressing B56δ mutant, p.E420K, a variant leading to intellectual disability in patients [20, 21]. Together, these data suggest B56δ-PP2A holoenzyme can regulate liprin-α1 dephosphorylation and LLPS that is impaired by a human mutation of intellectual disability.

## Results

### B56δ binds liprin-α1 via a short linear motif (SLiM)

We previously published that liprin-α1 binds B56δ [19]. In examining the peptide motifs on liprin-α1, we found 8 short linear interaction motifs (SLiM) LxxIxE for B56-PP2A family of holoenzymes [22, 23], with SLiM1, 4, 5 and 6 predicted to reside within disordered regions of liprin-α1 (Fig. 1A). While deletion of the first 30 N-terminal amino acids (encompassing SLiM1) did not affect B56δ binding (Fig. 1B), deletion of 152 or 180 amino acids of the liprin-α1 N-terminal region (additionally encompassing up to SLiM4) completely abolished B56δ binding (Fig. 1B). To determine the necessary SLiM for PP2A binding, we mutated critical residues (I/L/C and E) in the liprin-α1 SLIMs suspected to be functional within the N-terminal region (SLiMs 2, 3 and 4) and, as a negative control, within the C-terminal region (SLiMs 7 and 8) into valine (V) and/or alanine (A) (Fig. 1C). Only mutation of SLiM4 significantly affected B56δ binding (Fig. 1C). Since SLiM1 had previously been identified as a functional SLiM for binding to B56γ [16], we also verified its putative role in B56δ binding. To this end, SLiM1 and SLiM4 were mutated individually or together, tagged with GFP, and expressed in WT HEK293 cells (Fig. 1D). We found that B56δ binding to GFP-liprin-α1 SLiM1 mutant was unaffected, while its binding was significantly reduced when GFP-liprin-α1 contained the SLiM4 mutation, and was completely abolished when both SLiM1 and SLiM4 were mutated (Fig. 1D). Unbiased MS interactomics data of the liprin-α1 SLIM4 mutant (n=2) not only further confirmed decreased B56δ binding to this mutant, but also showed a concomitant decrease in binding of PP2A A and C subunits (Fig. 1E). Together, the data suggest SLiM4 is critical for B56δ-PP2A holoenzyme binding to liprin-α1.

**Figure 1:**
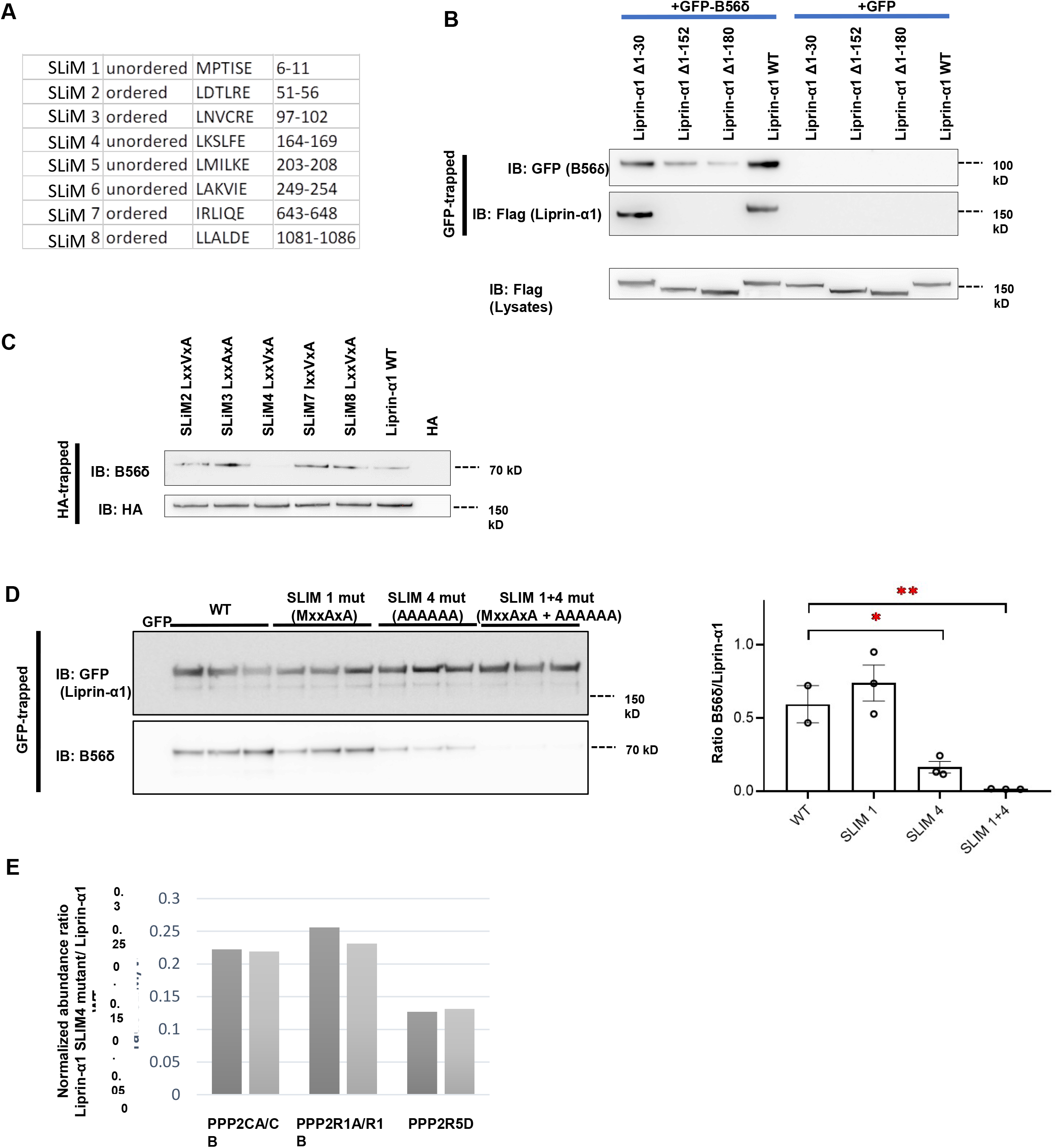
Identification of SLiM4 as the functional B56δ-interaction motif within liprin-α1. (A) Eight consensus LxxIxE short linear motifs (SLiMs) were identified in liprin-α1, potentially mediating the interaction with B56δ. The amino acid sequence and residue numbers are listed for each motif, as well as their presence within ordered versus disordered parts of the protein. (B) FLAG-tagged full-length liprin-α1 (WT) or N-terminal liprin-α1 deletions lacking the first 30 (Δ1-30), 152 (Δ1-152) or 180 amino acids (Δ1-180) were either co-transfected with GFP-tagged B56δ or GFP alone in HEK293T cells. Following GFP-trapping, the presence of FLAG-tagged proteins in the GFP pull downs was verified by immunoblotting. (C) HA-tagged full-length liprin-α1 (WT), indicated HA-tagged liprin-α1 SLiM mutants, or HA alone (control) were expressed in HEK293T cells, and binding of endogenous B56δ was assessed in anti-HA immunoprecipitates. (D) GFP-tagged full-length liprin-α1 (WT), GFP-tagged liprin-α1 SLiM1, SLIM4 or SLIM1+4 mutants, or GFP alone (control) were expressed in HEK293T cells, and binding of endogenous B56δ was assessed in GFP traps. Quantifications of the B56δ/liprin-α1 ratios in each condition are shown on the right (ANOVA, *: p<0.05, **:p<0.01), clearly demonstrating the functionality of SLiM4. (E) Effect of liprin-α1 SLiM4 mutation (LxxLxE to AAAAAA) on the presence of the PPP2R5D/PP2A holoenzyme in the liprin-α1 interactome, as determined by MS (n=2). Protein abundance of PP2A catalytic subunit (PPP2CA/CB), scaffolding subunit (PPP2R1) and regulatory B56 subunit (PPP2R5D) was normalized to liprin-α1 abundance in each condition, and for each independent experiment.

### Liprin-α1 mutant defective in B56**δ**-PP2A binding undergoes liquid-liquid phase separation

Liprin-α2 can undergo spontaneous formation of droplet-like structures and liprin-α3 undergo LLPS in response to PKC activation while liprin-α1 is not known to undergo spontaneous LLPS. In order to determine if phosphorylation of liprin-α1 can lead to its LLPS, we determined the effect of SLiM mutations on liprin-α1 LLPS. WT HEK293 cells were transfected with GFP-liprin-α1 with or without SLiM mutations. The SLiM4 mutation (and both SLiM1 and SLiM4 mutation) significantly increased GFP-liprin-α1 LLPS (Fig. 2A-B), suggesting B56δ targets PP2A to liprin-α1 to dephosphorylate residues involved in LLPS regulation. To determine that the droplets are indeed formed by LLPS, we subjected WT HEK293 cells containing GFP-liprin-α1 SLiM4 mutant droplets to fluorescence recovery after photobleaching (FRAP). LLPS droplets are dynamic and readily exchange with the surrounding cytosolic proteins, thus droplets should recover fluorescence over time. Droplets recovered to approximately 27% of the original fluorescence rapidly (t_1/2 max_ = 16.9s ± 8.09s) (Fig. 2C-E). We detected fusion events between GFP-liprin-α1 droplets as well, another hallmark of LLPS (Supplementary Fig. 1). Overall, we conclude B56δ-PP2A binding regulates liprin-α1 LLPS.

**Figure 2:**
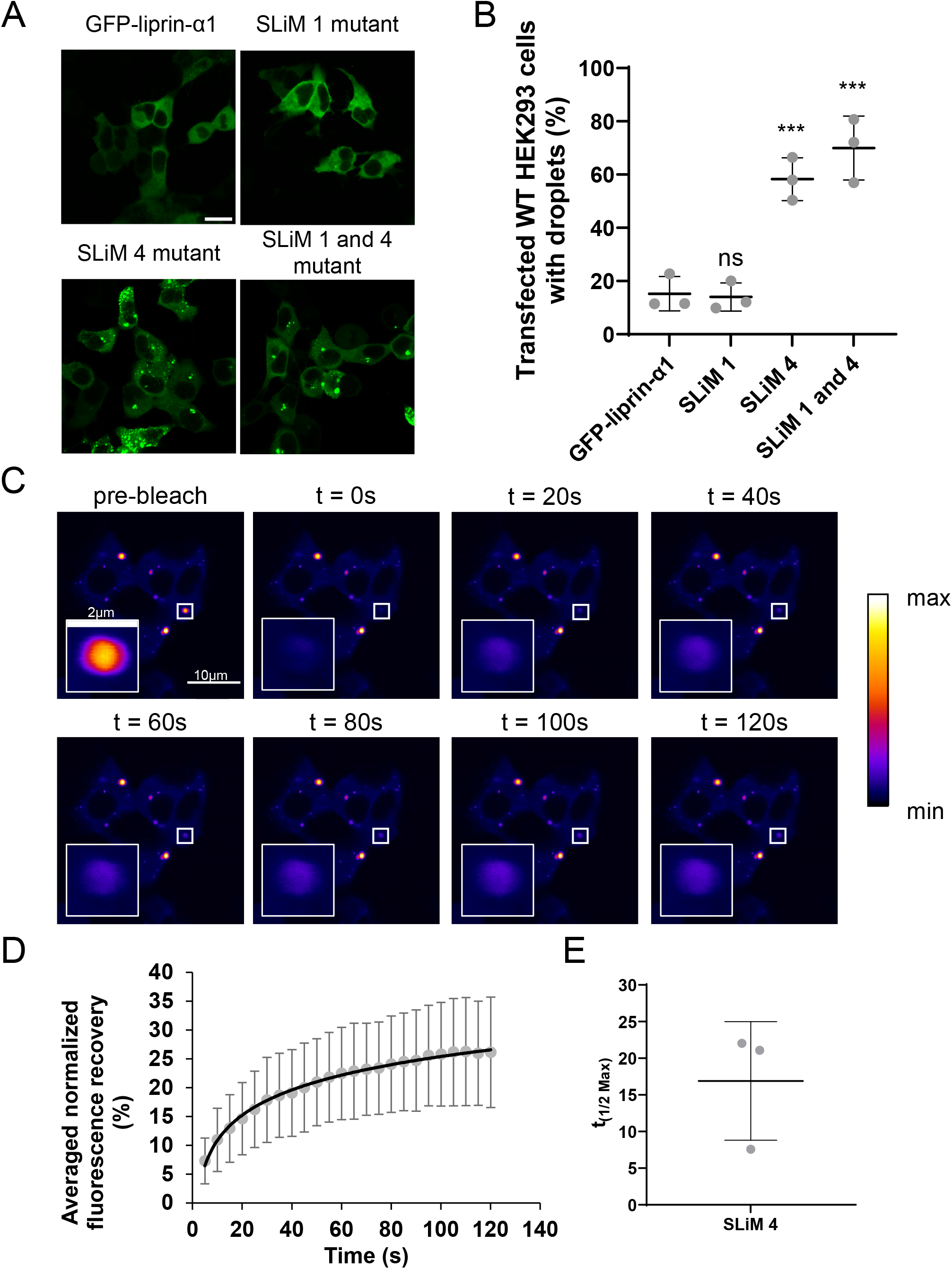
Liprin-α1 SLiM4 mutant significantly increases liprin-α1 liquid-liquid phase separation. (A) Example live confocal images of WT HEK293 cells transfected with GFP-liprin-α1, GFP-liprin-α1 SLiM 1 mutant, GFP-liprin-α1 SLiM 4 mutant, or GFP-liprin-α1 SLiM 1 and SLiM 4 mutant. Scale bar represents 20µm. (B) Quantification of the percentage of WT HEK293 cells with droplets. N=18 images/3 separate transfections (ANOVA with Tukey’s post hoc test, ns = not significant, ***p<0.001). (C) Example time-lapse confocal images of WT HEK293 cells transfected with GFP-liprin-α1 SLiM4 mutant. Droplets subject to fluorescence recovery after photobleaching (FRAP). 30 frames acquired with 5s between each frame. Bleaching occurred at frame 5 (t=0s). Images show the droplet directly before and after bleaching as well as the fluorescence recovery in 20s increments. (D) Normalized percentage of FRAP plotted over time. N=30 droplets/3 independent. Error bars indicate mean ± SD. (E) Quantification of the time to half of the maximal fluorescence recovery. Error bars indication mean ± SD.

### Mimicking liprin-α1 phosphorylation increased its liquid-liquid phase separation

Liprin-α1 is a well-known phosphoprotein based on phosphoSitePlus (https://www.phosphosite.org). Among the many phospho-sites listed for Liprin-α1 in PhosphoSitePlus, serine 763 (S763) residue is especially interesting because not only is it not conserved in other liprin-α isoforms (Fig. 3A), but it is also located in the intrinsic disordered region (IDR) which is usually important for regulating LLPS. We decided to examine the role of S763 in Liprin-α1 LLPS along with several other phospho-sites selected from PhosphoSitePlus (Fig. 3B). Of note, S1171 and S1172 phosphorylation were reported to increase in E420K phosphoproteome [24]. Phospho-mimetic mutations (serine (S) to glutamic acid (E)) were made at the following residues: S150, S666, S708, S763, and S1171/1172. WT HEK293 cells were transfected with GFP tagged liprin-α1 phospho-mimetic constructs and assessed for liprin-α1 LLPS (Fig. 4A-B). Phospho-mimetic mutation at S763 significantly increased GFP-liprin-α1 LLPS compared to the percentage of cells with LLPS when transfected with WT GFP-liprin-α1 (Fig. 4B). Our data thus suggest that serine 763 when phosphorylated can increase liprin-α1 LLPS.

**Figure 3:**
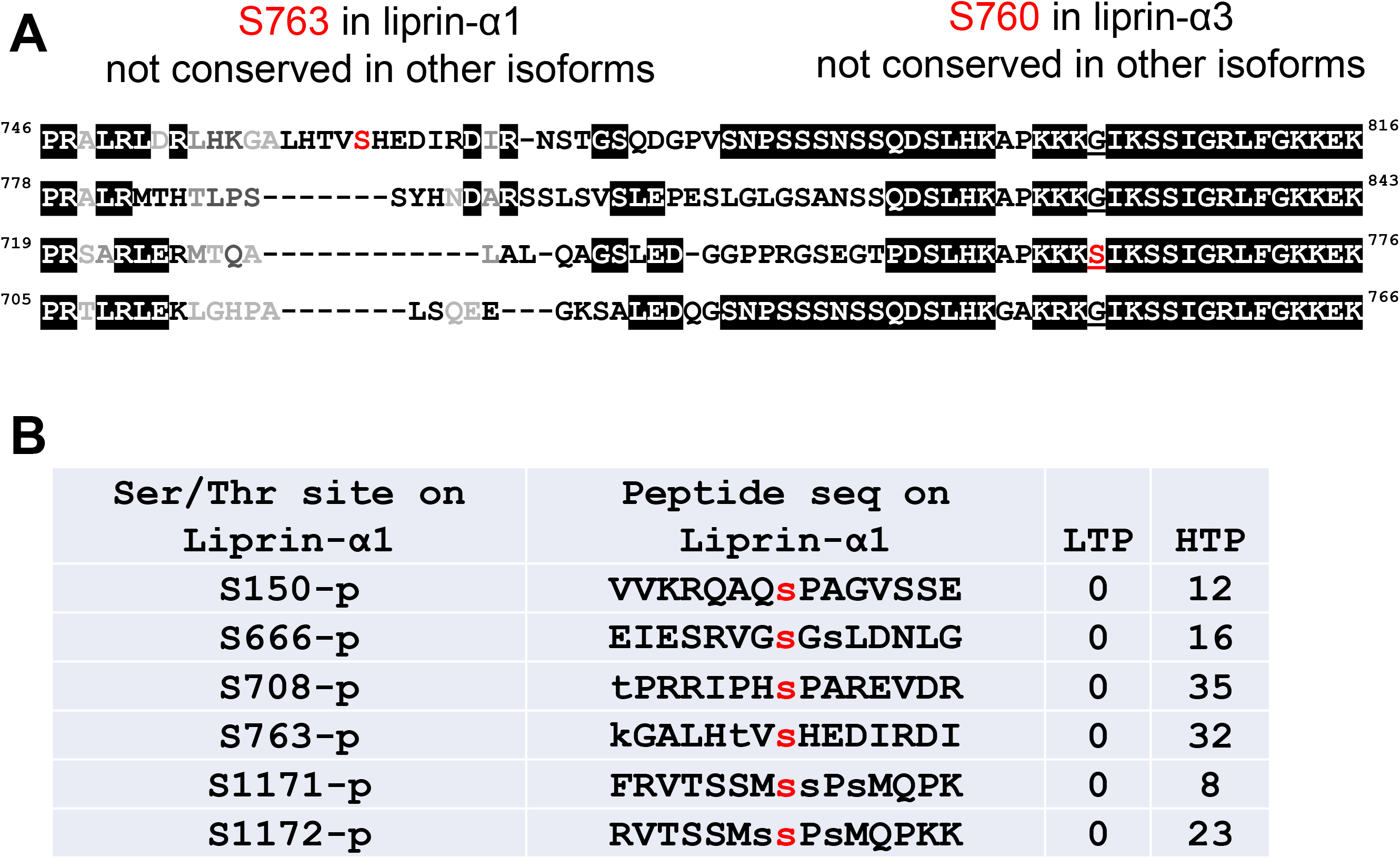
Potential phospho-sites in Liprin-α1 for LLPS studies. (A) alignment of Liprin-α1-4 sequences around S763 in Liprin-α1 (in red). Also shown is the position of S763 in Liprin-α3 (in red), which is also not conserved among Liprin-α isoforms. (B) List of phospho-sites in Liprin-α1 (in red) to be examined in this study. Also shown are the number of hits for the phospho-site in low through put (LTP) and high through put (HTP) studies (from PhosphoSitePlus: https://www.phosphosite.org).

**Figure 4.**
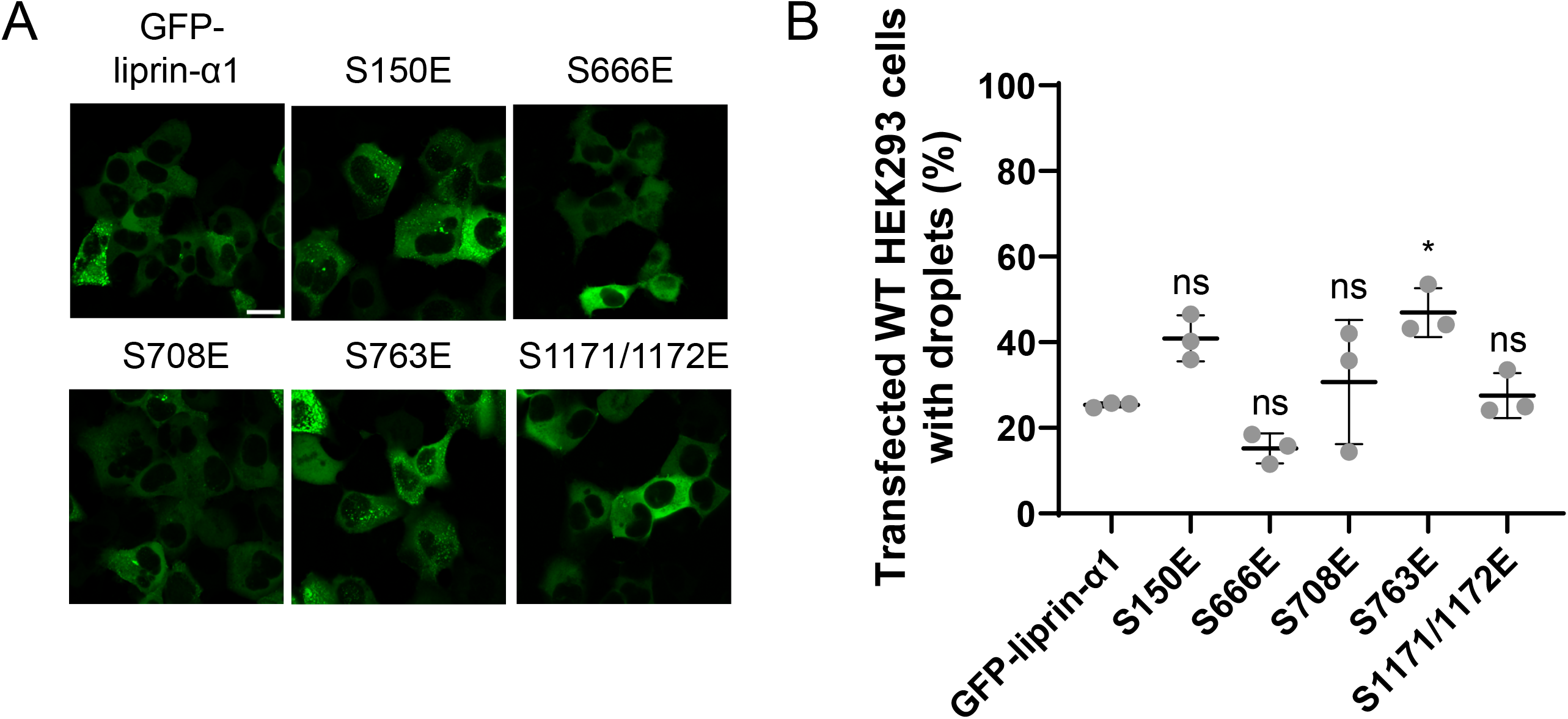
Mimicking liprin-α1 phosphorylation increased its liquid-liquid phase separation. (A) Example confocal images of live WT HEK293 cells transfected with GFP-liprin-α1 phospho-mimetic mutants (S150E, S666E, S708E, S763E, or S1171E/1172E). Scale bar indicates 20µm. (B) Quantification of the percentage of WT HEK293 cells transfected with phospho-mimetic GFP-liprin-α1 mutants with droplets. N=18 images/3 independent transfections. Error bars indication mean ± SD. (ANOVA with Tukey’s post hoc test, ns=not significant, *p<0.05, **p<0.01).

### Domains in liprin-α1 that contribute to its LLPS in B56δ KO HEK293 cells

Proteins that undergo LLPS often contain intrinsic disordered regions (IDR). We wanted to determine the contribution of IDR, along with other domains in liprin-α1, to its ability to undergo LLPS. Thus, we made GFP-liprin-α1 mutant constructs in which different domains were deleted: the IDR (ΔIDR), the three sterile alpha motifs (SAMs) (ΔSAMs), and the N-terminal dimerization region (amino acids 1-343 [25]: named ΔN) (Fig. 5A). For GFP-liprin-α1 (ΔN) constructs, we also mutated S763 to E mimicking phosphorylation (GFP-liprin-α1 (ΔN-S763E)).

**Figure 5:**
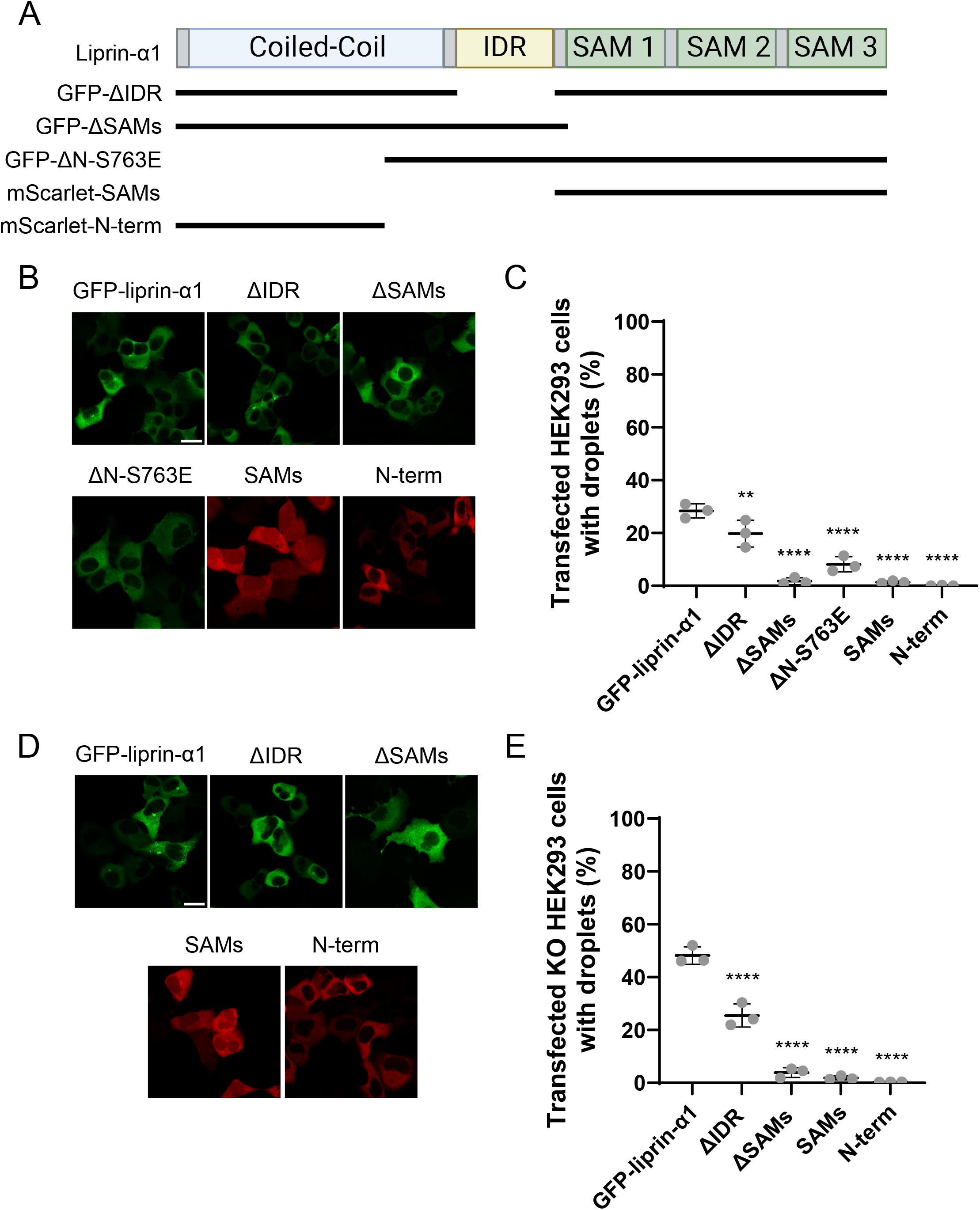
Domain mapping of liprin-α1 in its LLPS. (A) Schematic of liprin-α1 domains showing the design of constructs containing different liprin-α1 domain including deletion of the intrinsically disordered region (IDR, ΔIDR), deletion of all three SAM domains (ΔSAMs), deletion of the C-terminal dimerization domain (1-344) coupled with the S763E mutation (ΔN-S763E), and the SAM domains alone (SAMs), and the N-terminus only (1-344, N-term). Constructs are tagged with GFP or mScarlet as noted. (B) Example live confocal images of WT HEK293 cells transfected with GFP-ΔIDR, GFP-ΔSAMs, GFP-ΔN-S763E, mScarlet-SAMs, or mScarlet-N-term. Scale bar indicated 20µm. (C) Quantification of the percentage of WT HEK293 cells transfected with various liprin-α1 domain mutants containing droplets. N=18 images/3 independent transfections. Error bars indication mean ± SD. (ANOVA with Tukey’s post hoc test, **p<0.01, ****p<0.0001) (D) Example live confocal images of B56δ KO HEK293 cells transfected with GFP-ΔIDR, GFP-ΔSAMs, mScarlet-SAMs, or mScarlet-N-term. Scale bar indicated 20µm. (E) Quantification of the percentage of KO HEK293 cells transfected with various liprin-α1 domain mutants containing droplets. N=18 images/3 independent transfections. Error bars indication mean ± SD. (ANOVA with Tukey’s post hoc test, ****p<0.0001).

Additionally, we made two mScarlet tagged liprin-α1 constructs. The first contains only the 3 SAMs (mScarlet-SAMs) to determine whether the SAMs domain can promote LLPS on its own as they are supramodules for structural and functional complex formation [26] (Fig. 5A). The second mScarlet construct contains the N-terminus (amino acids 1-343; named N-term) (Fig. 5A).

Because we found KO HEK293 cells had increased GFP-liprin-α1 LLPS (Fig. 4B-C), we wanted to assess the necessity of liprin-α1 domains in this system. Expression of GFP-ΔIDR significantly reduced droplet formation in KO HEK293 cells near what was observed in WT HEK293 cells, however, expression of GFP-ΔSAM decreased droplet formation even further (Fig. 5D-E). Lastly, we observed that deletion of N-terminal dimerization domain also significantly decreased GFP-liprin-α1 droplet formation (Fig. 5D-E). Our data suggest full-length liprin-α1 is necessary for its LLPS in KO HEK293 cells. We also performed LLPS experiments for GFP-liprin-α1 domain-deletion mutant constructs in WT HEK293 cells (Fig. 5B-C). GFP-ΔIDR resulted in a low level of droplets, similar to WT HEK293 cells expressing GFP-liprin-α1 (Fig. 5C). GFP-ΔSAMs expressing WT HEK293 cells resulted in nearly no cells with droplets, suggesting again the critical role of SAM domains in droplet formation. Deletion of the first 343 amino acids (ΔN-S763E) also resulted in very little droplet formation even with the phospho-mimetic mutations, suggesting dimerization is critical for LLPS of liprin-α1 (Fig.5C).

### B56δ potently inhibits liprin-α1 liquid-liquid phase separation

Liprin-α1 SLiM4 mutations significantly decreased B56δ binding and resulted in increased phase separation in WT HEK293 cells. To further assess the contribution of B56δ to liprin-α1 phase separation, we generated a B56δ knock-out HEK293 (KO) cell line by using a CRISPR editing system. Our immunostaining indicated B56δ is successfully knocked out (Fig. 6A). Interestingly expression of GFP-liprin-α1 WT in KO HEK293 cells led to significant droplet formation (Fig. 6B and C). The data suggest B56δ may be the primary B subunit targeting PP2A to liprin-α1, dephosphorylating it and suppressing its LLPS.

**Figure 6:**
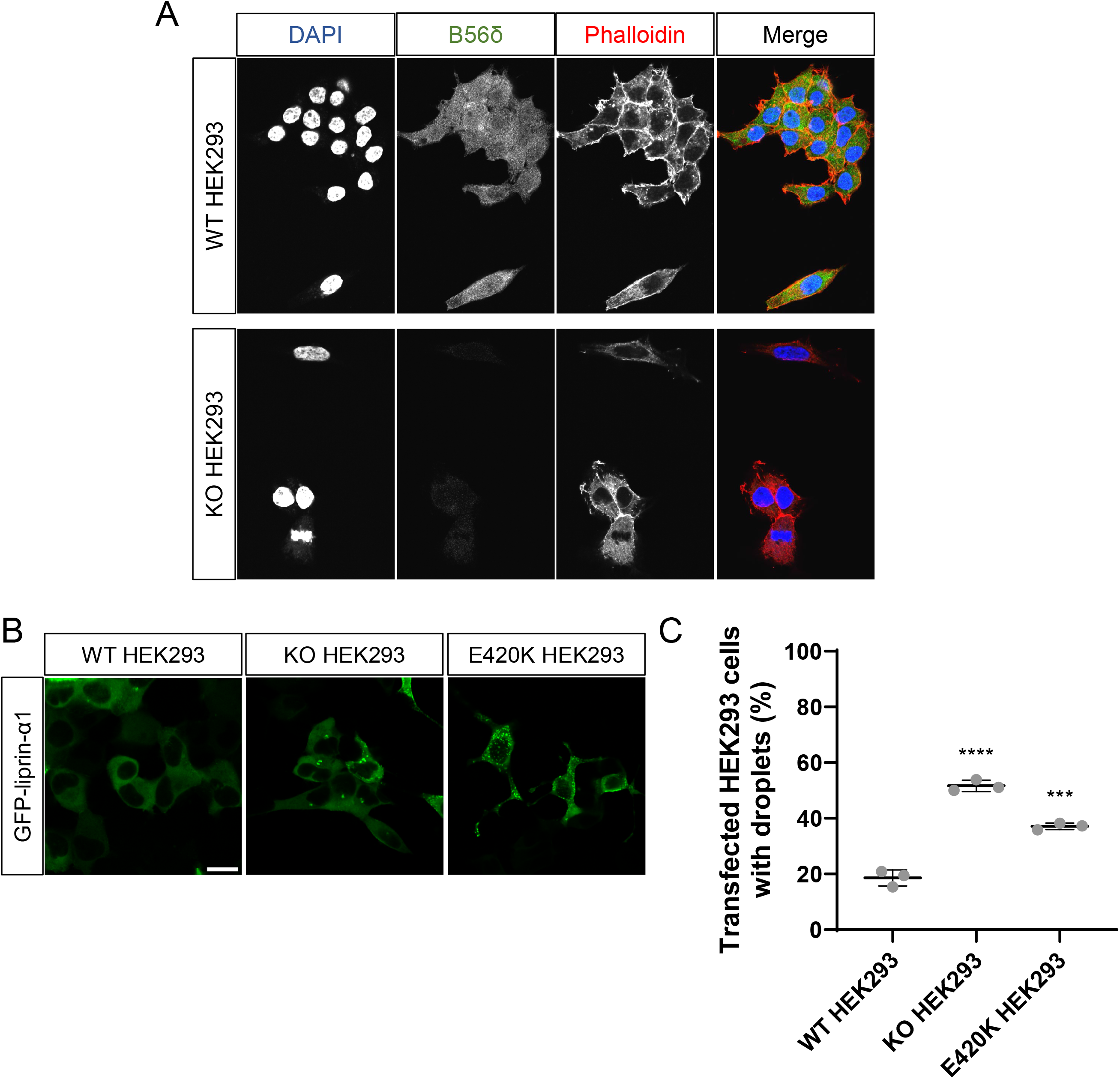
Increased liprin-α1 LLPS in B56δ knock-out (KO) HEK293 cells and human de novo mutation E420K homozygous HEK293 cells. (A) Immunohistochemistry validation of B56δ knock-out HEK293 cells. (B) Representative confocal images of WT HEK293, B56δ KO HEK293 cells, or E420K homozygous HEK293 cells transfected with GFP-liprin-α1. Scale bar indicates 20µm. (C) Quantification of the percentage of cells containing droplets. N=18-36 images/3 independent transfections (ANOVA with Tukey’s post hoc test, ***p<0.001, ****p<0.0001). Error bars indicate mean ± SD.

We next designed liprin-α1 phospho-null constructs corresponding to the phospho-mimetic mutants used previously. Phospho-null mutants were transfected in KO HEK293 cells, but no singular phospho-blocking mutant or combination of mutants tested were able to decrease droplet formation (Supplementary Fig. 2). We concluded that B56δ-PP2A holoenzyme could access additional phosphorylation sites, potently inhibiting liprin-α1 LLPS.

### B56δ human *de novo* mutation E420K impairs B56δ-PP2A’s suppressing effect on liprin-α1 liquid-liquid phase separation

Human *de novo* mutations in B56δ were found to disrupt its binding to liprin-α1 [19], therefore we wanted to explore the impact on liprin-α1 phase separation. We used isogenic homozygous HEK293 cell lines with single genomic base at E420 in both alleles of B56δ gene mutated to K using a CRISPR single base editor [24] to assess the impact of this human mutation on GFP-liprin-α1 driven phase separation. E420K homozygous HEK293 cells showed significantly increased GFP-liprin-α1 driven phase separation, suggesting mutant B56δ-PP2A is not able to properly dephosphorylate liprin-α1 (Fig. 6B and C).

## Discussion

LLPS is a major mechanism of concentrating proteins in nano-domains in cells for efficient and specific spatial and temporal signaling. While many studies focus on the regulation of LLPS via kinases, we identified a phosphatase responsible for negatively regulating the LLPS of liprin-α1. Our experiments revealed B56δ targets PP2A to liprin-α1 via binding to SLiM4 and mutating SLiM4 significantly promoted GFP-liprin-α1 LLPS. Through information from PhosphoSitePlus, literature and educated guess, we studied several liprin-α1 phosphorylation sites. A phospho-mimetic mutant at S763 was sufficient to increase liprin-α1 LLPS. We also revealed the expression of the pathogenic HJS1 p.E420K variant increased GFP-liprin-α1 LLPS.

Many studies identified proteins highly concentrated in nano-domains such as the postsynaptic density and focal adhesions that undergo LLPS. PSD95 is a prime example of postsynaptic protein that can undergo LLPS spontaneously and with other synapse proteins including SynGAP [4, 27], TARP γ8/stargazin [4, 6], NMDA receptors, and other synapse scaffolding proteins such as GKAP, Shank and Homer [3]. Inhibitory synaptic scaffolding protein gephyrin [28] can also undergo LLPS with GABAa receptors. GIT1 and β-PIX are both synapse and focal adhesion proteins. GIT1 can undergo LLPS in a concentration dependent manner [7]. β-PIX cannot undergo LLPS on its own, it can facilitate GIT1 LLPS [7]. Similarly, focal adhesion molecule paxillin can further facilitate GIT1-β-PIX LLPS [7]. However, whether phosphorylation can regulate the LLPS for these proteins are not known. Among these synaptic proteins, phosphorylation of SynGAP by CaMKII abolishes the interaction between SynGAP with PSD95 [29], suggesting that phosphorylation could potentially affect LLPS between the two proteins. Many proteins localized in the active zone in the presynaptic terminal can also undergo LLPS [1, 5, 30–32]. Among them liprin-α3 undergoes LLPS in response to PKC activation [1], however, the identity of opposing phosphatase in liprin-α3 LLPS regulation is not known. Another well-known example of phosphorylation regulated LLPS is related to tau. Tau undergoes spontaneous LLPS and phosphorylation of tau has been initially identified as the main driver for this process [33]. However, once again, no studies examined the contribution of phosphatases regulating tau LLPS. The abundant evidence for protein LLPS is contrasted with little known about phosphatase regulation in this important cellular process.

B56δ targets PP2A catalytic subunit to liprin-α1 via direct interaction between B56δ and liprin-α1 and between B56δ and PP2A [19]. Our study mapped the interaction between liprin-α1 and B56δ: B56δ specifically binds to SLiM4 on liprin-α1. Phospho-mimetic mutant S763E significantly increased GFP-liprin-α1 LLPS; however, S150E, S666E, S708E, and S1172E/1172E were not sufficient for increasing GFP-liprin-α1 LLPS. S763 is located in the intrinsic disordered region (IDR) on liprin-α1. Interestingly, S763 is unique to liprin-α1. This is reminiscent of S760 on IDR of liprin-α3 which is not present in other liprin-α isoforms and is sufficient for liprin-α3 LLPS in response to PKC activation. Our data and studies of liprin-α3 suggest unique regulation of different mammalian liprin-α isoforms

Liprin-α homologue in *Caenorhabditis elegans*, SYD-2 also undergoes LLPS using a distinct mechanism. SAD-1 kinase activation promotes SYD-2 LLPS [5]. Interestingly, mutant SYD-2 with deletion of C-terminal SAM domains undergo spontaneous LLPS. The SAM domains interact with the N-terminus of SYD-2 to prevent LLPS [5]. SAD-1 kinase phosphorylates three sites on the SAM1 motif and relieves the inhibition of SYD-2 LLPS [5]. In our study, we did not observe spontaneous LLPS for the analogous deletion of liprin-α1 (GFP-liprin-α1 (ΔSAMs)). This is also true for liprin-α3 [1], suggesting divergent mechanism of mammalian liprin-α LLPS regulation from *Caenorhabditis elegans* homologue.

Our observation of spontaneous liprin-α1 LLPS in B56δ KO cells definitively determined that B56δ contributes to the dephosphorylation of liprin-α1 resulting in suppression of its LLPS. LxxIxE motif on liprin-α1 is a SLiM recognized by B56 family proteins. Our SLiM4 mutant of liprin-α1 does not bind to B56δ. S763A, the phosphorylation blocking mutant, cannot decrease the spontaneous GFP-liprin-α1 LLPS in B56δ KO HEK293 cells. This suggests B56δ could potentially access another or multiple phosphorylation sites on liprin-α1 whose phosphorylation will increase in B56δ KO cells and are also sufficient to drive LLPS. This scenario would indicate there is more than one liprin-α1 phosphorylation event triggering LLPS and that B56δ-PP2A can dephosphorylate all these phosphorylation sites, potently suppressing liprin-α1 LLPS. A single mutation in B56δ, E420K, leading to intellectual disability also resulted in decreased binding to liprin-α1 [19]. Consistent with this finding, we found significant LLPS in HEK293 cells with PPP2R5D containing the E420K mutation. Our data suggest a single mutation in B56δ can disrupt its regulation of liprin-α1 LLPS.

Our domain deletion of liprin-α1 indicated full-length protein is required for LLPS. Specifically, the N-terminal dimerization domain, middle intrinsic disordered regions, and the C-terminal SAM domains are all important for liprin-α1 LLPS. Our data are consistent with what was reported for liprin-α3 [1]. Yet, our findings contrast with other protein examples, such as PSD95, SynGAP, and GIT1. In these examples, critical domains can be isolated and undergo LLPS spontaneously. In phosphorylation dependent LLPS, it is conceivable that phosphorylation sites may reside between critical domains for inducing protein conformation change, and/or in domains that affect protein-protein interaction critical for LLPS, thus requiring full-length or near full-length protein.

In summary, we found that liprin-α1 undergoes phosphorylation-promoted LLPS. Moreover, we have identified that B56δ binds liprin-α1 via SLiM4, putatively accessing serine 763 on liprin-α1 to potently suppress its ability to form biocondensates.

## Experimental Methods

### DNA constructs

The liprin-α1 cDNA was obtained from DNASU (clone HsCD00044464), and subcloned into different eukaryotic expression vectors by a PCR-based approach (proofreading PWO polymerase): pMB001 (N-terminal HA-tag), pCMV-3xFLAG (N-terminal FLAG-tag) and pEGFP-C1 (N-terminal EGFP-tag). All constructs were expressed under the CMV promoter. Deletion mutants were generated by PCR (PWO polymerase) and appropriate primers, and cloned into pCMV-3xFLAG. GFP-liprin-α1 SLiM mutants, phospho-mimetic, phospho-blocking, and domain deletion mutants were made using site directed mutagenesis (NEBasechanger or NEBuilder (New England Biolabs (MA)). Relevant G-blocks for NEBuilder were ordered from Twist Bioscience (South San Francisco, CA). All constructs were verified by Sanger sequencing.

### Immunoprecipitation and Western Blotting

B56δ-liprin-α1 binding was assessed by various methodologies. Initially, plasmids encoding FLAG-tagged liprin-α1 and GFP-tagged B56δ were co-transfected in HEK293T cells (3µg of each, using PEI-max), and soluble lysates were subjected to anti-GFP pulldown, essentially as described [19, 34] Following SDS-PAGE and immunoblotting, the pulldowns were counterstained with anti-GFP (Cell Signaling Technologies, # 2555S) and anti-FLAG antibodies (Sigma, # F1804). In addition, HA-tagged or GFP-tagged liprinα1 or its variants were expressed in HEK293T cells, isolated by anti-HA immunoprecipitation [19, 34, 35] or GFP trapping [19] and counterstained for HA (Sigma, # H9658) and endogenous B56δ (Abcam, # ab188323), or GFP and endogenous B56δ, respectively. Western blotting and quantification of bands were performed as previously described [20].

### Quantification of droplets in HEK293 cells

HEK293 cells were plated onto culture dishes and transfected with 2ug of the plasmid of interest. DNA constructs used for any co-transfections were added in 1:1 ratio. Live cells were imaged approximately 18 to 21 hours post transfection. Cells were kept in tissue culture medium without phenol red. Images were acquired using a Keyence microscope with a 40x objective or an Olympus confocal microscope using a water-immersion 25X objective. Quantification was completed manually to determine the total number of transfected cells and the percentage of the transfected cells that contained spherical droplets. The experimenter was blind to the conditions using an image randomizer plug-in (Fiji) for data analyses.

### Fluorescence recovery after photobleaching

FRAP was completed using live HEK293 cells plated on 60mm culture dishes. 18 to 21 hours after transfection, cells were transferred to the confocal microscope for experiment. Single droplets were bleached individually by drawing a region of interest around the droplet. Images were acquired every 5 seconds for a total of 30 cycles. Each droplet was bleached at cycle 5 with 488 nm laser wavelength for 2 seconds. The fluorescence intensity for each droplet was normalized and averaged before being plotted over time. The start of recovery until maximal recovery was plotted and fit to a logarithmic curve. The time to half maximum recovery (t_1/2 max_) was calculated. For assessment of fusion evets, time-lapse imaging was performed. Fusion events were defined as two droplets merging into a larger droplet.

### Liprin-α1 interactomics

GFP-Liprin-α1 WT and SLiM4 variant were overexpressed in HEK293T cells, lysed in NET buffer (50 mM Tris.HCl pH 7.4, 150 mM NaCl, 15 mM EDTA and 1% Nonidet P-40) supplemented with cOmpleteTM protease inhibitor and PhosSTOP^TM^ phosphatase inhibitor cocktails (Roche). Lysates were subsequently subjected to GFP trapping, washing and unbiased interactomics essentially as described in [36]. Protein abundances were obtained by averaging the abundance of the 3 most abundant peptides per protein.

### Immunocytochemistry

Immunocytochemistry was performed as in [37]. In detail, HEK293 cells were plated on poly-d-lysine coated #1.0 glass coverslips. Cells were fixed with 4% paraformaldehyde for 15 min, permeabilized with 0.2% Triton X-100 in PBS for 10 min and blocked in a PBS based solution containing 1%BSA/5%FBS/0.1% Triton X-100 for 1 hour. Coverslips were incubated with primary antibody (rabbit anti-PPP2R5D, abcam188323, 1:250) diluted in blocking buffer overnight at 4°C. The following day, after wash with PBS, coverslips were incubated with Alexa-conjugated secondary antibody (goat anti-rabbit Alexa Fluor 488, 1:1000) in blocking buffer for 1 hour. Cells were also labeled with phalloidin conjugated to Alexa Fluor 594 in blocking buffer according to manufacturer protocol (Thermofisher A30107). After incubation with DAPI for 5 min and subsequent PBS wash, coverslips were mounted with Prolong Diamond Anti-Fade Mount. Coverslips were imaged using Olympus confocal microscope equipped with a 60X oil immersion objective. Identical settings were used to capture images between cell lines.

### Statistics

GraphPad Prism (10.1.2) was used for statistical analysis and data visualization. For liquid-liquid phase separation experiments, statistical significance was assessed using one-way ANOVA testing. Equality of group variances was tested with the Brown-Forsythe test before performing a one-way ANOVA. Tukey *post hoc* comparisons were performed for one-way ANOVAs. The mean and standard deviation are displayed in all figures.

## Acknowledgements

This work was supported by NIH R01MH128279 and the URMC Startup fund to HX, and the Jordan’s Guardian Angel Foundation fund (University of California Reagents) to HX, VJ, BEW and REH. IV is a junior postdoctoral research fellow of the FWO-Vlaanderen. The authors declare no competing financial interests. BEW is co-founder of Turkey Creek Biotechnology LLC (Waverly, TN).

## Supplemental Figure Legends

**Supplemental Figure 1:**
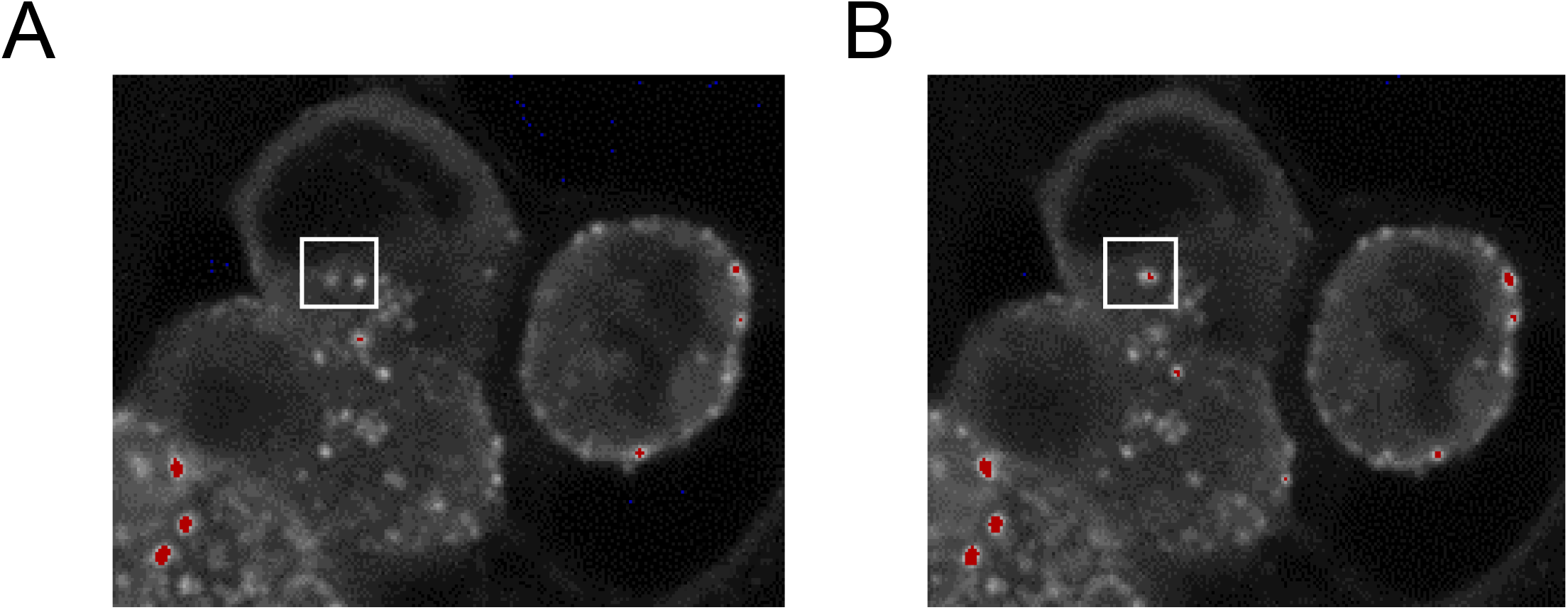
GFP-liprin-α1 SLiM 4 mutant droplets are dynamic and undergo fusion events. Live confocal timelapse imaging of WT HEK293 cells transfected with GFP-liprin-α1 SLiM 4 mutant. (A) Before fusion. (B) Droplets after fusion event.

**Supplemental Figure 2:**
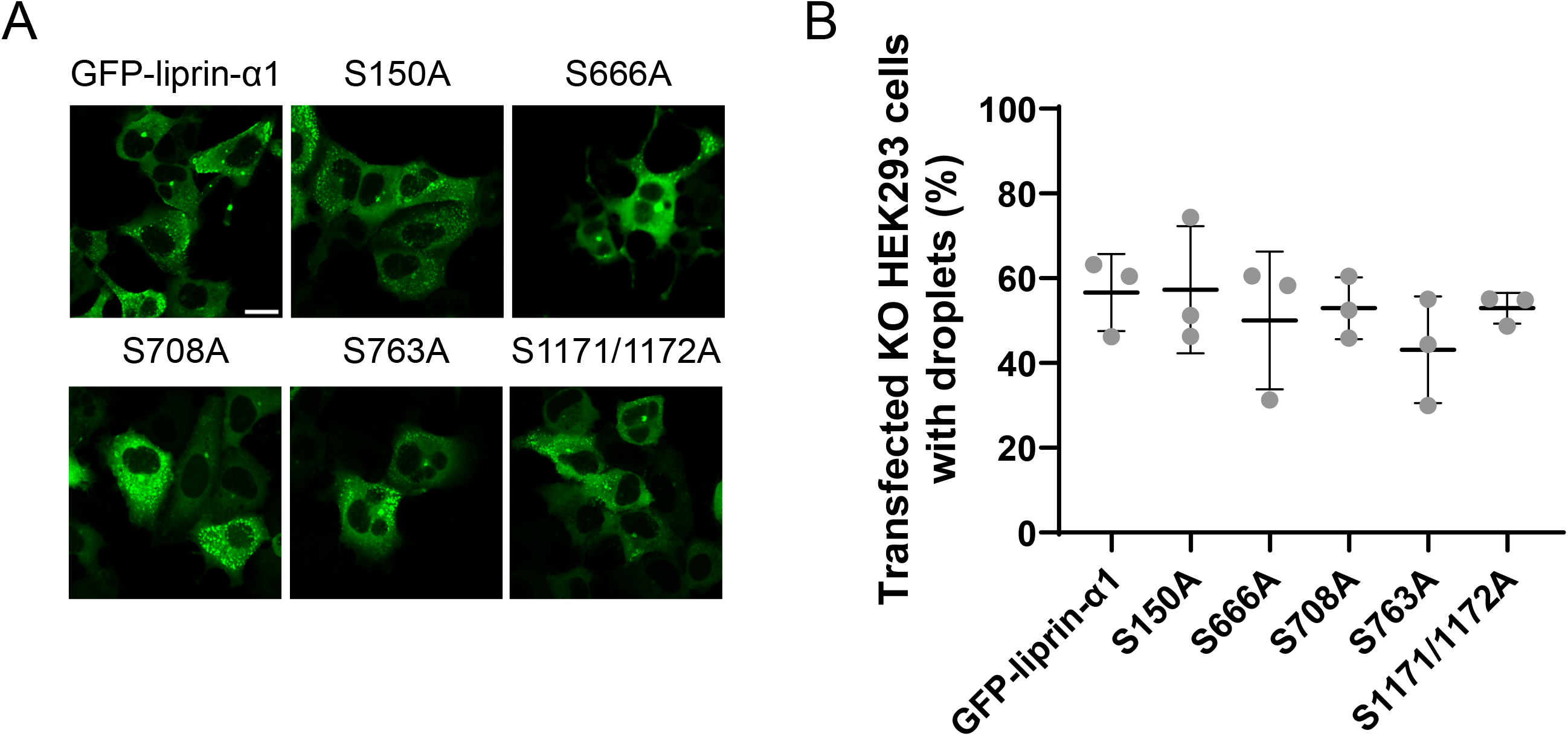
individual or double phospho-null liprin-α1 mutants do not block LLPS in KO HEK 293 cells. (A) Representative confocal images of live B56δ KO HEK293 cells transfected with GFP-liprin-α1 phospho-null mutants (S150A, S666A, S708A, S763A, and S1171a/S1172A). Scale bar indicates 20µm. (C) Quantification of the percentage of KO HEK293 cells transfected with phospho-null GFP-liprin-α1 mutants with droplets. N=18 images/3 independent transfections. (ANOVA with Tukey’s post hoc test). Error bars indication mean ± SD.

